# A nanoscale atlas of extracellular vesicles and particles in *Drosophila* olfactory sensilla

**DOI:** 10.64898/2026.05.30.728996

**Authors:** Pawel Vijayakumar, Kalyani Cauwenberghs, Muskaan Garg, Jonathan Choy, Shadi Charara, Quintyn McKaughan, Chih-Ying Su

## Abstract

Extracellular particles, including non-vesicular extracellular particles (NVEPs) and extracellular vesicles (EVs), are emerging as key contributors to sensory signaling, yet their ultrastructural organization within native tissues remains underexplored. Native tissues preserve extracellular particle organization and heterogeneity that are often lost during dissociation. Using cryofixation-based serial block-face scanning electron microscopy, we generated a nanoscale atlas of NVEPs and EVs across 352 *Drosophila* antennal olfactory sensilla (∼70% coverage). Segmentation of more than 7,800 extracellular particles revealed distinct populations differing in morphology, size, electron density, and sensillum-class distribution. Analyses of EV biogenesis identified multivesicular bodies and putative membrane budding in auxiliary cells. Furthermore, sensilla housing degenerating neurons exhibited marked accumulation of EVs and NVEPs, accompanied by increased auxiliary-cell EV biogenesis. These findings provide a large-scale ultrastructural characterization of extracellular particles in native sensory tissues and establish a foundation for understanding how extracellular particles are distributed, generated, and function during sensory signaling and degeneration.

## INTRODUCTION

Extracellular particles, including extracellular vesicles (EVs) and non-vesicular extracellular particles (NVEPs), are increasingly recognized as important mediators of intercellular communication in the nervous system (Remans et al., 2025; Van Niel et al., 2018; Welsh et al., 2024; Yáñez-Mó et al., 2015). EVs can transport proteins, lipids, nucleic acids, and signaling molecules between cells, and EV-mediated signaling has been linked to neuron–glia communication, synaptic modulation, and neurodegeneration (Bavisotto et al., 2019; Ikezu et al., 2024; Pistono et al., 2021). Notably in sensory systems, certain ciliated neurons in *C. elegans* actively release EVs which mediate both local signaling within sensory organs and animal-to-animal communication (Wang et al., 2014, 2020, 2021). Yet despite growing interest, relatively little is known about the ultrastructural organization of extracellular particles within native sensory tissues. This knowledge gap persists in part because most studies rely on dissociated or purified preparations that disrupt original tissue architecture, or on light microscopy approaches that lack the spatial resolution necessary to resolve nanoscale structural heterogeneity. Characterizing these ultrastructural features will provide insights into how extracellular particles are distributed, generated, and function within native sensory microenvironments.

The *Drosophila* olfactory system is well suited for bridging this knowledge gap because extracellular interactions occur within individual sensory hairs (sensilla) that contain olfactory neurons and associated auxiliary cells within a confined, lymph-filled microenvironment (Kaissling, 1986). Fly olfactory sensilla fall into four morphological classes—coeloconic, basiconic, intermediate, and trichoid—in ascending order of length (Nava Gonzales et al., 2021). Olfactory receptor neurons (ORNs) across these classes also differ in functional tuning. Trichoid ORNs primarily detect pheromones, basiconic and intermediate ORNs respond to fruit-related volatiles, and coeloconic ORNs are generally tuned to acids and amines (Dweck et al., 2015; Hallem & Carlson, 2006; Lin et al., 2016; Silbering et al., 2011; van der Goes van Naters & Carlson, 2007; Yao et al., 2005). Together, this structural and functional organization partitions odor detection into specialized compartments at the periphery.

In *Drosophila*, each sensillum typically contains two and up to four ORNs. The pairings of ORNs within individual sensilla are stereotyped (Benton et al., 2009, 2025; Couto et al., 2005; Fishilevich & Vosshall, 2005; Hallem et al., 2004) and are largely conserved across *Drosophila* species (Auer et al., 2020; Linz et al., 2013; Prieto-Godino et al., 2017; Stensmyr et al., 2003), suggesting that compartmentalization of ORNs within sensilla is functionally important. Indeed, neighboring ORNs interact within the sensillum via ephaptic inhibition to compute odor valence, allowing each sensillum to operate as a peripheral processing unit (Ng et al., 2020; Puri et al., 2024; Su et al., 2012; Wu et al., 2022; Zhang et al., 2019). These findings highlight the need for tightly coordinated development and maintenance of compartmentalized ORNs to ensure proper functional coupling within individual processing units.

In addition to ORNs, each olfactory sensillum contains three types of auxiliary cells—thecogen, trichogen, and tormogen cells (Figure 1—figure supplement 1)—arranged in a nested, cup-like organization (Nava Gonzales et al., 2021; Shanbhag et al., 2000). The innermost thecogen cell ensheathes parts of the ORN somata, as well as the inner dendrites and proximal outer dendrites. Surrounding the thecogen cell is the trichogen cell, which extends extensive microlamellae into the sensillum lumen. The outermost tormogen cell partially encloses the trichogen cell near the cuticle base and similarly projects microlamellae into the lumen (Choy et al., 2025; Nava Gonzales et al., 2021). All sensillum classes contain one auxiliary cell of each type, except coeloconic sensilla, which contain an additional trichogen cell (Nava Gonzales et al., 2021). Auxiliary cells support olfactory function by maintaining transepithelial potential, and secreting odorant-binding proteins and odorant-degrading enzymes into the sensillum lymph (Kaissling, 1986; Larter et al., 2016; Leal, 2013; J. S. Sun et al., 2018). As shared cellular components, these auxiliary cells are ideally positioned to coordinate the development and function of compartmentalized ORNs.

Indeed, EVs released from auxiliary cells likely contribute to intercellular communication. For example, Osiris 8 (Osi8), a transmembrane protein expressed in the tormogen cells of trichoid sensilla, is detectable in the sensillum lumen as vesicle-like puncta, suggesting that it is released via EVs. Loss of Osi8 markedly reduces pheromone responses in both Or47b and Or88a ORNs co-housed in the at4 trichoid sensillum (Scalzotto et al., 2022). Consistent with these findings, our previous volume electron microscopy (EM) study identified EVs that appeared to bud from tormogen cell membranes (Nava Gonzales et al., 2021), highlighting a potential role for EV-mediated signaling in trichoid sensilla.

In parallel, NVEPs may represent an additional and underexplored form of intercellular communication within the sensillum. Unlike EVs, NVEPs lack a surrounding lipid membrane and consist of protein aggregates, lipoprotein-like particles, or other macromolecular assemblies in the extracellular space (Welsh et al., 2024). These particles could provide an alternative means for molecular exchange or signaling between cells within a sensillum. However, their presence, abundance, and morphological features within olfactory sensilla remain largely unknown because NVEPs are generally smaller than EVs (often within the tens-of-nanometers range) (Shahi et al., 2024), making detection difficult with conventional light microscopy.

Beyond EVs and NVEPs released from living cells, apoptotic bodies generated during programmed cell death likely serve as an additional source of intercellular communication (Battistelli & Falcieri, 2021). Consistent with reported ORN apoptosis in the antenna (Chihara et al., 2014; Fernández-Hernández et al., 2020; Takeuchi et al., 2022), a small fraction of sensilla in our published EM volumes (0.4%) lacked ORNs, with or without intact auxiliary cells (Nava Gonzales et al., 2021), indicating that neuronal loss had occurred, and that our SBEM volumes likely captured ORNs undergoing degeneration. Thus, our EM volumes provide a unique opportunity to determine whether sensilla containing degenerating ORNs exhibit altered EV or NVEP abundance, and to examine their ultrastructural features and biogenesis.

Our previous volume EM analysis revealed the presence of EVs in multiple sensillum classes, but whether EVs are differentially distributed or enriched in specific sensilla remains unknown. Their nanoscale morphological diversity has also not been systematically characterized. EV size provides clues to their biogenesis: small vesicles (30–150 nm) are typically exosomes derived from intracellular multivesicular bodies (MVBs), whereas larger vesicles likely correspond to microvesicles (100–1000 nm) or apoptotic bodies (1000–5000 nm) that bud or bleb from the plasma membrane. In addition to size, the luminal electron density of EVs likely indicates cargo abundance: electron-dense vesicles are often associated with higher concentrations of biomolecular cargo (e.g., nucleic acids and proteins), whereas electron-lucent vesicles may reflect more dilute or less structured contents (Doyle & Wang, 2019; Malkin & Bratman, 2020; Shahi et al., 2024; Singh et al., 2024; Welsh et al., 2024). Characterizing the distribution of EV sizes and luminal densities across sensillum classes may uncover class-specific organizational features linked to the distinct functions or physiological properties of individual olfactory processing units.

Here, we examine serial block-face scanning electron microscopy (SBEM) volumes of the *Drosophila* antenna generated using cryofixation-based methods (Denk & Horstmann, 2004; Tsang et al., 2018). Cryofixation by high-pressure freezing followed by freeze substitution faithfully preserves cuticle-encased sensory tissues and enables accurate nanoscale morphological characterization not achievable with conventional chemical fixation (Shanbhag et al., 1999, 2000, 2001; Tsang et al., 2018). By providing high-quality ultrastructural preservation, this approach enables cross-sensillum surveys of EVs and NVEPs, nanoscale morphometric analyses, and identification of their cellular origins. Together, these data establish a detailed nanoscale atlas of EVs and NVEPs in the olfactory periphery, which thus provides a foundation for future studies into their biogenesis, cargo, and function during sensory signaling and degeneration.

## RESULTS

Our laboratory previously generated multiple antennal SBEM volumes (Choy et al., 2025; Nava Gonzales et al., 2021; Zhang et al., 2019), each acquired from a subregion of the antenna in a different fly. One such volume spans a substantial portion of the antenna, encompassing approximately 70% of all antennal olfactory sensilla (Choy et al., 2025). We focused our analysis on this large volume to systematically compare EV/NVEP distribution and morphology within a single animal. To assess the generality of our observations, we further examined EVs and NVEPs from three smaller SBEM volumes, each sampling a different antennal subregion (Figure 1A).

**Figure 1.**
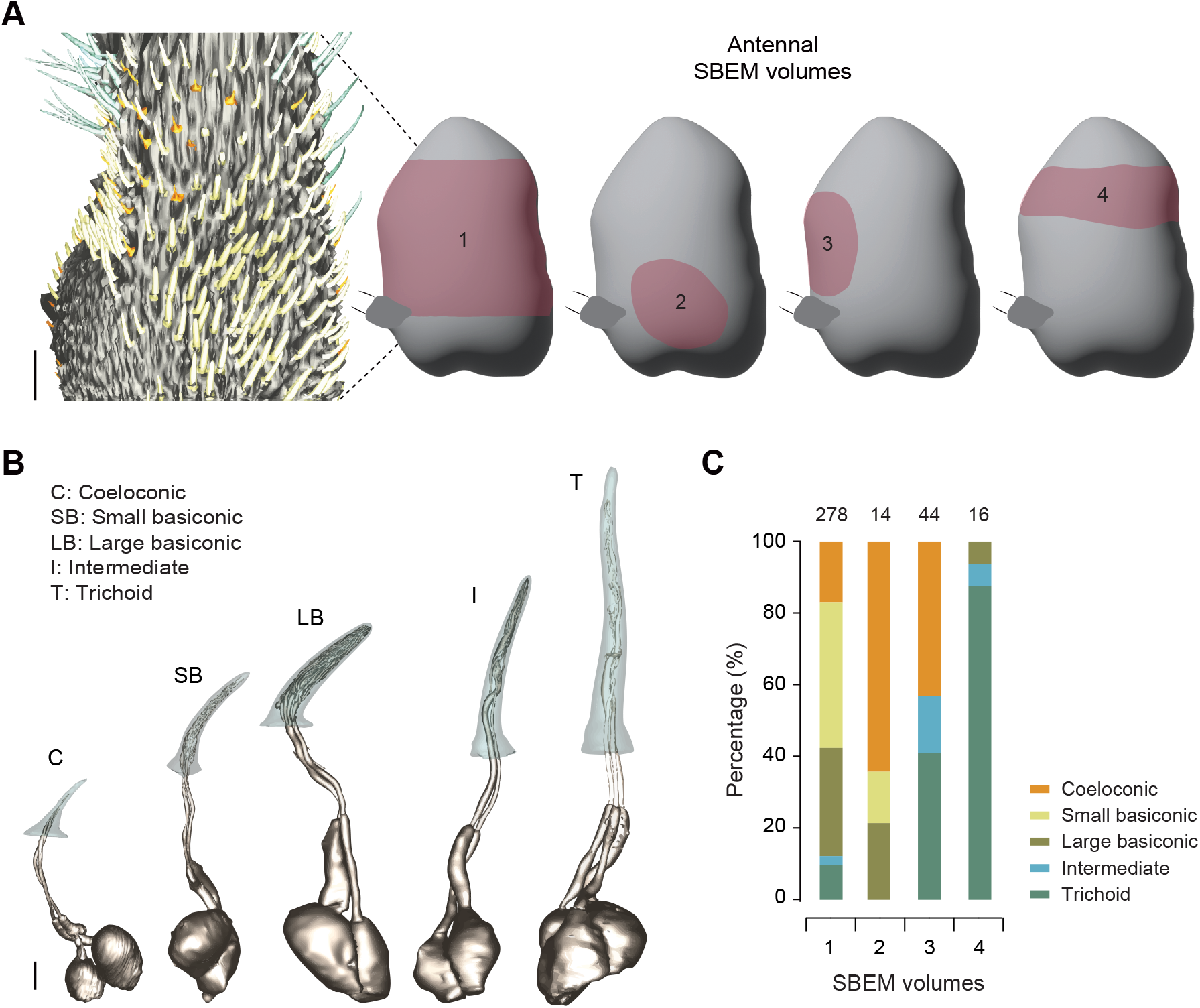
Different sensillum classes and their representation across SBEM volumes. **(A)** Schematic of antennal regions sampled in each SBEM volume analyzed in this study (pink-shaded areas). Inset: the largest volume, showing 3D reconstructions of the antenna and sensillum cuticles for basiconic (yellow), coeloconic (orange), intermediate (blue), and trichoid (teal) sensilla. Scale bar, 15 µm. **(B)** Representative 3D models of the major morphological classes of olfactory sensilla and their associated ORNs. Scale bar, 2 µm. **(C)** Proportional representation of sensillum classes across SBEM volumes. In volumes 2–4, only a subset of sensilla within each volume was sampled and analyzed. The total number of analyzed sensilla is indicated above each bar.

Collectively, these SBEM volumes covered all major morphological classes of olfactory sensilla—coeloconic, basiconic, intermediate, and trichoid. Basiconic sensilla were further subdivided into small and large types (Figure 1B). Consistent with the spatially segregated (rather than salt-and-pepper) organization of sensillum classes across the antenna (de Bruyne et al., 2001; Grabe et al., 2016; Shanbhag et al., 1999), their representation varied across volumes (Figure 1C). Each morphological class was sampled in at least two independent SBEM volumes.

Across these datasets, we analyzed 352 sensilla, representing ∼70% of antennal sensilla (Shanbhag et al., 1999), selected based on image quality and completeness, including only sensilla fully contained within the SBEM volumes. Among these, 32 sensilla housed degenerating ORNs. In total, we segmented 4,793 EVs and 3,030 NVEPs for morphological and morphometric analyses. Together, these datasets enabled a systematic survey of EVs and NVEPs, facilitating comparisons of their prevalence, abundance, morphology, and distribution under both normal and degenerating conditions.

### Morphological comparison of EVs and NVEPs

From the four SBEM volumes, we analyzed 320 sensilla with intact ORNs and no apparent signs of degeneration, including 105 small basiconic, 81 large basiconic, 69 coeloconic, 51 trichoid, and 14 intermediate sensilla. The unequal sampling across classes reflects both their relative abundance in the antenna (basiconic > trichoid > coeloconic > intermediate) (Shanbhag et al., 1999) and their representation within the volumes. From these sensilla, 3,311 extracellular particles (including both EVs and NVEPs) within the sensillum lumen were segmented. Based on ultrastructural morphology, 1,397 particles were enclosed by a clearly defined membrane and were classified as EVs (Figure 2A), whereas 1,914 particles lacked a discernible bounding membrane and exhibited high electron density throughout the particle, and were classified as NVEPs (Figure 2B) (Welsh et al., 2024).

**Figure 2.**
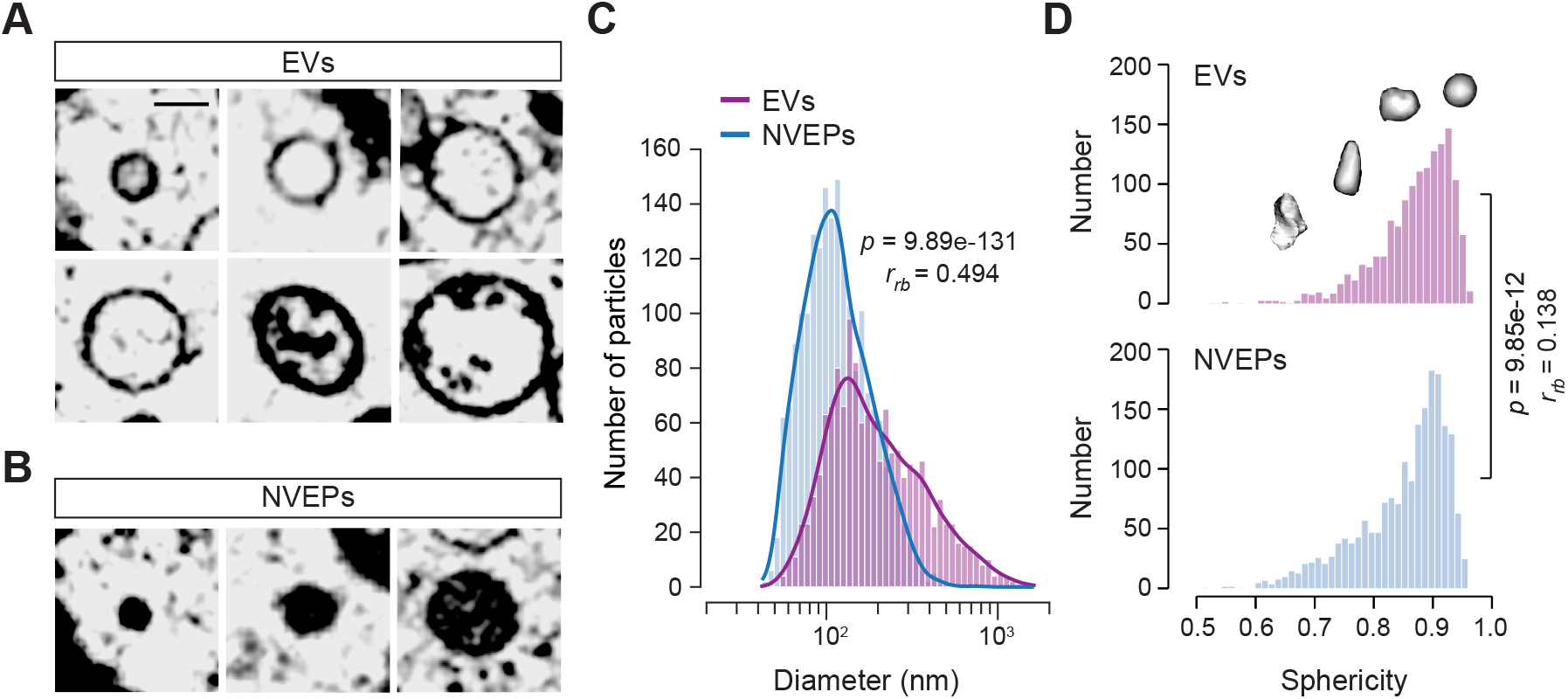
Morphological comparison of EVs and NVEPs (A–B) Representative SBEM images of EVs (**A**) and NVEPs (**B**), arranged in ascending order of diameter. The scale bar shown in the first panel applies to all images. Scale bar, 250 nm. **(C)** Distributions of EV and NVEP diameters. Curves represent kernel density estimates. Median diameters are 177 nm for EVs and 111 nm for NVEPs. Group differences were assessed using the Mann-Whitney U test, and effect sizes were quantified using the rank-biserial correlation (*r_rb_*) (*n* = 1,397 EVs and 1,914 NVEPs). Hartigan’s dip test supported unimodality for overall EV (*p* = 0.990) and NVEP distributions (*p* = 0.981). **(D)** Distributions of EV and NVEP sphericity, with representative 3D models illustrating increasing sphericity from left to right. Median sphericity values are 0.88 for EVs and 0.87 for NVEPs. Statistical analysis as in (**C**).

To compare particle size, we estimated an effective diameter for each particle based on its volume and analyzed the resulting size distributions. Although particle size was not used as a classification criterion, NVEPs were generally smaller than EVs, with median diameters of 111 nm and 177 nm, respectively (Figure 2C). Notably, the size distributions overlapped in the lower diameter range, indicating that small particles are present in both populations. However, EVs exhibited a broader distribution with a pronounced rightward tail, reflecting a higher prevalence of larger particles compared to NVEPs. Collectively, EVs were significantly larger than NVEPs.

We next quantified particle shape using sphericity, defined as the ratio of the surface area of a sphere with the same volume to the particle’s actual surface area. Under this definition, a perfect sphere has a sphericity of 1, whereas increasing shape irregularity results in lower values. Both EVs and NVEPs exhibited high sphericity with median values of 0.88 and 0.87, respectively, indicating that most particles are nearly spherical (Figure 2D). The high sphericity of both populations indicates that cryofixation preserved these nanoscale particles in a near-native hydrated state, avoiding the flattened “cup-shaped” artifacts commonly associated with conventional chemical fixation (Raposo & Stoorvogel, 2013; Welsh et al., 2024).

### Comparison of EVs and NVEPs across sensillum classes

#### Relative abundance of EVs and NVEPs

Both EVs and NVEPs were detected in all sensillum classes (Figure 3A). To assess whether each class exhibited a bias toward one particle type, we analyzed the EV ratio, defined as the number of EVs divided by the total number of EVs and NVEPs. A value of 0.5 indicates equal representation of EVs and NVEPs.

**Figure 3.**
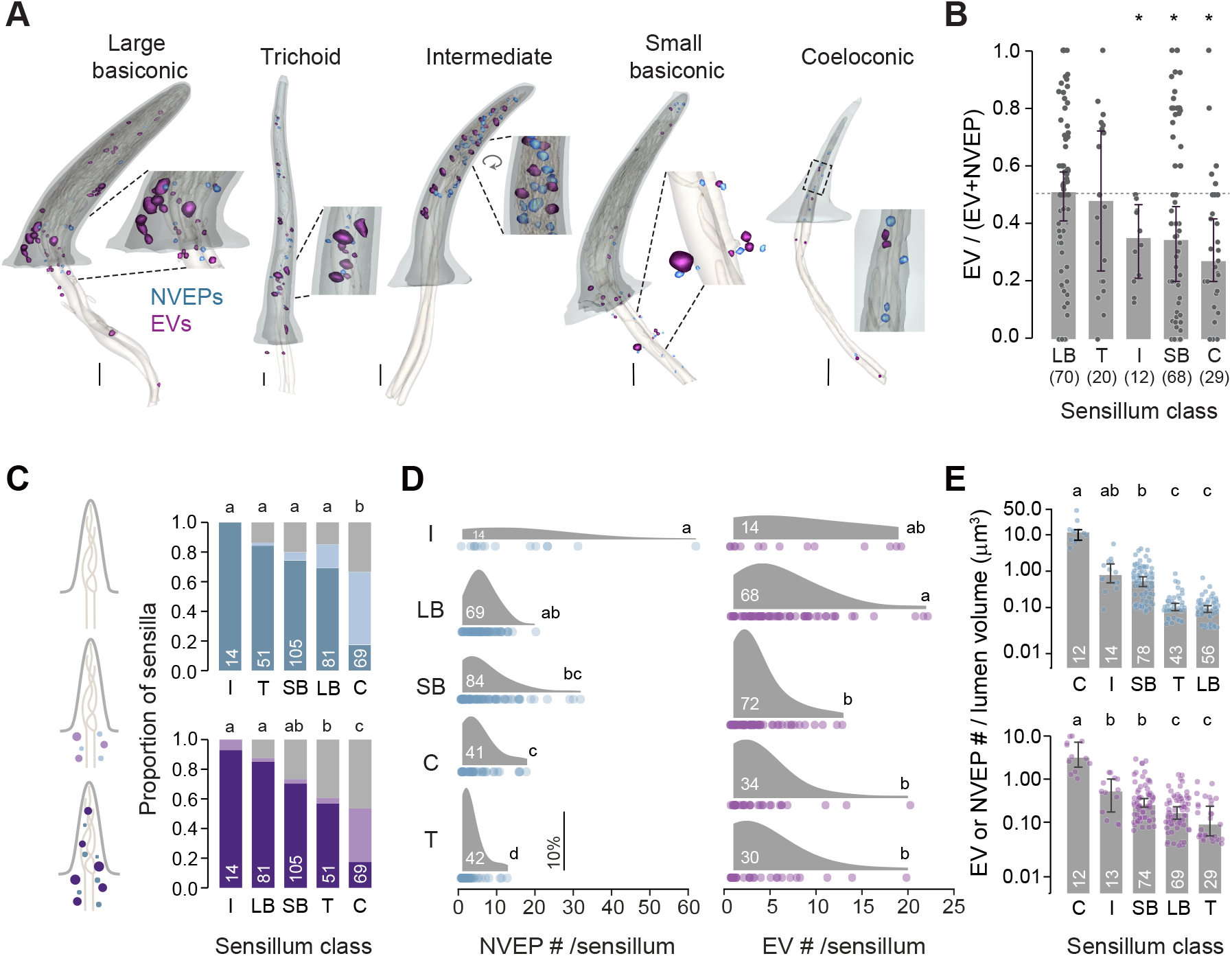
Comparison of EVs and NVEPs across sensillum classes. **(A)** Representative 3D models of major olfactory sensillum classes, showing sensillum cuticle, lumen, ORN outer dendrites, EVs (purple), and NVEPs (blue). Insets: magnified views. Scale bar, 1 µm. **(B)** EV and NVEP prevalence. The EV ratio is defined as the number of EVs divided by the total number of EVs and NVEPs. Each datapoint represents one sensillum; only sensilla with ≥5 particles were considered. Numbers of analyzed sensilla are indicated. Median ± 95% CI. Dashed line indicates an EV ratio of 0.5. Significance was assessed by Wilcoxon signed-rank test against 0.5 (\**p* < 0.05). **(C)** EV and NVEP distribution. Left: Schematic illustrating three categories of particle localization: 1) no EVs/NVEPs (gray); 2) EVs/NVEPs restricted to the sensillum lumen below the cuticle base (light shade); and 3) EVs/NVEPs present in the lumen both above and below the cuticle base (dark shade). Right: Stacked bar graphs across sensillum classes. Different letters indicate statistically significant differences between groups (Chi-square test for independence with FDR-corrected Chi-square *post hoc* tests, *p* < 0.05). **(D)** EV and NVEP number. Only sensilla containing at least one EV or NVEP were considered and outliers were removed using Tukey’s extreme fences. Distributions are represented as kernel density estimates with individual data points shown along the x-axis. Kruskal–Wallis test followed by FDR-corrected Dunn’s post hoc tests, *p* < 0.05. Scale bar, 10% density. **(E)** Volume-normalized EV and NVEP abundance. Only sensilla containing at least one EV or NVEP above the cuticle base were included. Each datapoint represents a single sensillum; median ± 95% CI. Statistical analysis as in (**D**).

The median EV ratios of large basiconic and trichoid sensilla did not differ significantly from 0.5, whereas intermediate, small basiconic, and coeloconic sensilla showed median ratios significantly below 0.5, indicating a population-level NVEP bias (Figure 3B). Nevertheless, EV ratios varied considerably among sensilla of the same class, and distributions overlapped broadly across sensillum classes. Consequently, no pairwise differences between classes survived FDR-corrected Dunn’s *post hoc* comparisons despite a trend across classes (Kruskal–Wallis test, *p* = 4.76 × 10^-2^). Thus, while NVEPs were generally more abundant than EVs at the population level in most sensillum classes, both particle types co-occurred in individual sensilla with highly variable relative proportions.

#### EV and NVEP distribution

The position of extracellular particles relative to the cuticle base is likely to inform their cellular targets and sites of action, particularly in the lumen above the cuticle base, where ORN outer dendrites reside. We therefore examined particle position distribution by plotting each particle’s relative distance from the sensillum base (Figure 3—figure supplement 1). This continuous distribution analysis revealed distinct, though variable, class-specific spatial patterns.

To summarize these distributions, we additionally classified sensilla into three categories: no particles, particles present only below the cuticle base, or particles present above the base (including those also with particles below, as sensilla with particles exclusively above the base were rare) (Figure 3C). Most sensilla contained EVs and/or NVEPs, while few were entirely particle-free. Of note, coeloconic sensilla—the smallest sensillum type (Figure 1B)—showed the highest proportion entirely lacking particles, and when present, EVs and NVEPs were largely restricted below the cuticle base (Figure 3C; Figure 3—figure supplement 1) Thus, while most sensilla harbored particles above the base, coeloconic sensilla showed either no particles or particles restricted below the base, suggesting class-specific differences in sites of action.

#### EV and NVEP number

To compare particle abundance across sensillum classes, we counted EVs and NVEPs within the sensillum lumen. These snapshot counts reflect the steady-state pool at fixation; they do not capture dynamic rates of production or turnover. Quantification of particle number per sensillum revealed class-dependent differences for NVEPs. Among sensilla containing at least one particle, the median number was 4 NVEPs per sensillum. NVEP counts varied markedly across classes: intermediate sensilla showed the broadest distribution and the highest median (Mdn = 8.5), followed by large basiconic (Mdn = 6), small basiconic (Mdn = 4.5), and coeloconic sensilla (Mdn = 4), whereas trichoid sensilla contained the fewest NVEPs (Mdn = 2). These differences were statistically significant overall, with intermediate sensilla differing from lower-count classes and trichoid sensilla exhibiting the lowest NVEP numbers (Figure 3D, left).

EV counts followed a similar pattern but with a narrower distribution (Figure 3D, right). Intermediate and large basiconic sensilla showed broader ranges and higher counts (both Mdn = 5), whereas small basiconic and coeloconic sensilla were lower (both Mdn = 3), and trichoid sensilla again contained the fewest EVs (Mdn = 2). The overall median was 3 EVs per sensillum, with relatively modest and largely non-significant differences between sensillum classes compared to those observed for NVEPs.

In summary, extracellular particle abundance varies by sensillum type, whereby intermediate and large basiconic sensilla generally contain more particles, while trichoid sensilla contain the fewest.

#### Volume-normalized EV and NVEP abundance

Sensillum size varied markedly across classes (Figure 1B). If EV and NVEP production rates were comparable across sensillum classes, one would expect differences in lumen size alone to account for variation in total particle number. To test whether particle counts scale with sensillum size, we compared EV and NVEP numbers normalized to lumen volume. For this analysis, we considered sensilla containing at least one particle above the cuticle base, as lumen volume was quantified only for this region (see Materials and Methods for details). Volume estimation below the cuticle base was not feasible due to the complex microlamellar structures of surrounding auxiliary cells (Nava Gonzales et al., 2021). Sensillum lumen sizes differed substantially across classes, with large basiconic sensilla exhibiting the highest volumes (29.92 ± 0.52 µm^3^, mean ± SEM), followed by trichoid (23.53 ± 0.48 µm^3^), intermediate (11.78 ± 0.64 µm^3^), and small basiconic sensilla (9.61 ± 0.19 µm^3^), while coeloconic sensilla were much smaller (0.56 ± 0.03 µm^3^) (see Source Data for Figure 3).

If lumen volume were the primary determinant of particle abundance, volume-normalized particle abundance should be roughly uniform across all sensillum types. However, this was not observed. Smaller sensilla (e.g., coeloconic) exhibited substantially higher particle concentrations than larger sensilla (e.g. trichoid or large basiconic) (Figure 3E). This inverse relationship indicates that particle abundance is not a simple function of available lumen space. Rather, it suggests that underlying factors—such as differential production, release, or clearance rates—are specific to each sensillum class.

### Variation in EV properties across sensillum classes

Among the 1,397 EVs described in the previous section, ultrastructural analysis revealed three categories based on intraluminal electron density: dense (*n* = 730), lucent (*n* = 421), and cargo-filled (*n* = 246) (Figure 4A). These are morphological categories strictly based on ultrastructure, not verified biological classes. Dense and lucent EVs, both displaying uniform intraluminal density, were defined relative to the surrounding sensillum lymph: dense EV interiors were similar to or more electron-dense than the lymph, whereas lucent EV interiors were less electron-dense than the surrounding lymph. In contrast, cargo-filled EVs showed heterogeneous intraluminal density: a light interior with discrete, electron-dense inclusions. These categories were readily distinguishable in SBEM images and observed across sensillum classes. They likely reflect differences in luminal content: dense and cargo-filled EVs appeared more macromolecule-rich than lucent EVs.

**Figure 4.**
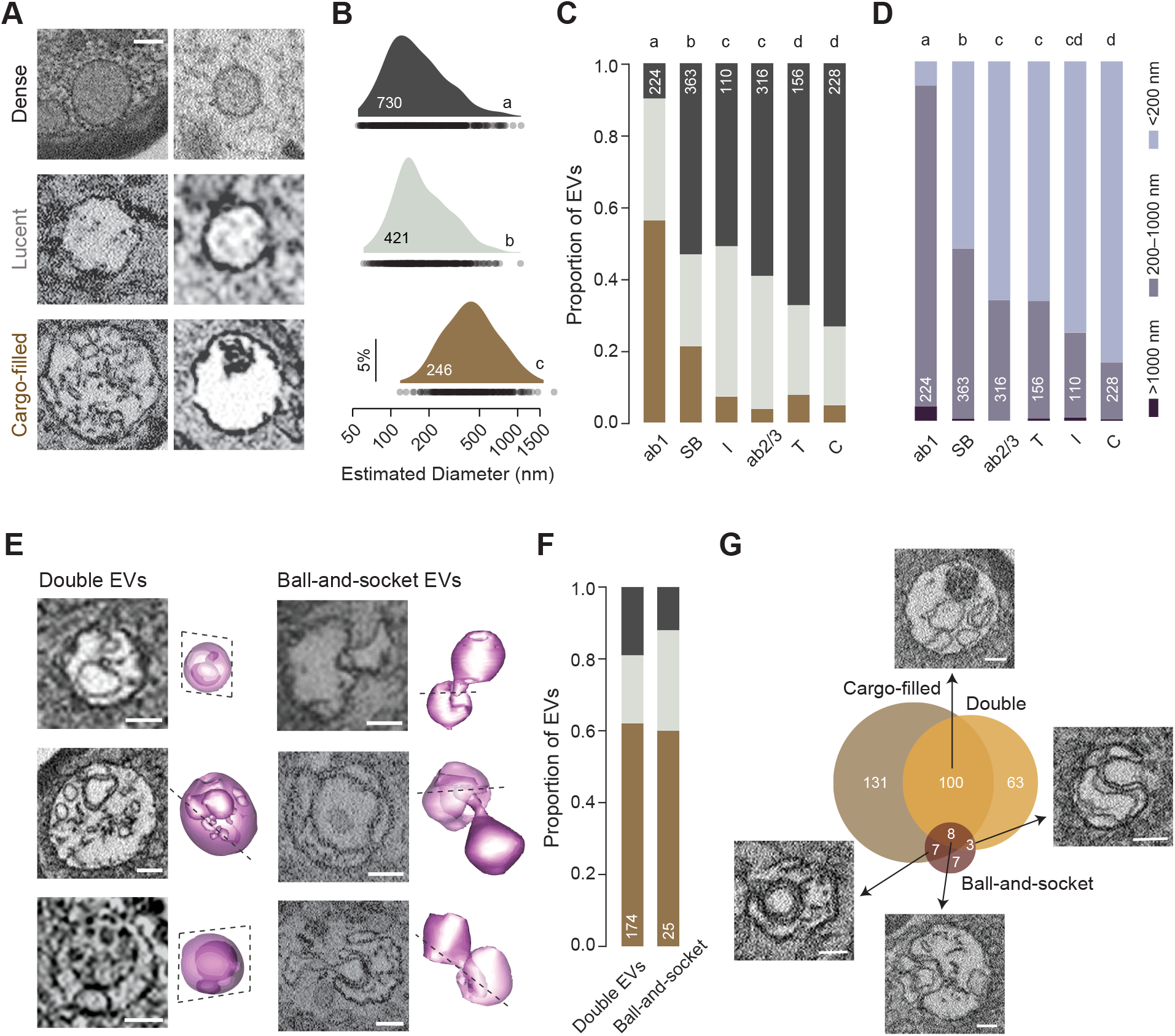
Variation in EV properties across sensillum classes. **(A)** Representative SBEM images illustrating three categories of EV intraluminal density: dense, lucent, and cargo-filled. Two independent examples are shown for each category. Scale bar, 100 nm (applies to all panels). **(B)** Size distributions of EVs by intraluminal density category, shown as kernel density estimates with individual data points plotted along the x-axis. Sample sizes for each category are indicated. Different letters denote statistically significant differences between groups (Kruskal–Wallis test with FDR-corrected Dunn’s *post hoc* comparisons, *p* < 0.05). Hartigan’s dip test supported unimodality for dense (*p* = 0.993), lucent (*p* = 0.993), and cargo-filled EVs (*p* = 0.951). **(C)** Stacked bar plots showing the proportion of EV density categories across sensillum classes. Numbers of sensilla analyzed per class are indicated. Sensilla lacking EVs in the lumen were excluded from the analysis. Sensillum classes: large basiconic (ab1, ab2/3), small basiconic (SB), intermediate (I), trichoid (T), and coeloconic (C). Chi-square test for independence with FDR-corrected Chi-square *post hoc* tests, *p* < 0.05. **(D)** Stacked bar plots showing the proportion of EV diameter categories across sensillum classes. Categories represent the largest vesicle observed in a sensillum; EVs exceeding 200 nm are consistent with a microvesicle-like size range. Statistical analysis as in (**C**). **(E)** Representative SBEM images and corresponding 3D models of EV structural variants: double EVs (left) and ball-and-socket EVs (right). Scale bars, 250 nm. **(F)** Stacked bar plots showing the distribution of intraluminal density categories within double EVs and ball-and-socket EVs, highlighting enrichment of the cargo-filled category. **(G)** Venn diagram showing overlap between cargo-filled EVs, double EVs, and ball-and-socket EVs. Numbers indicate counts of EVs in each category and their intersections. Representative SBEM images of EVs from individual overlapping subsets are shown. Scale bars, 250 nm.

We next asked whether these density classes differ in size. EV diameter distributions varied significantly across categories (Figure 4B). Cargo-filled EVs were substantially larger and displayed a broader, right-skewed distribution (median diameter = 440 nm), whereas dense EVs were the smallest (147 nm), and lucent EVs showed intermediate sizes (161 nm), indicating that intraluminal density is associated with distinct EV sizes.

How are EV density categories distributed across sensillum classes? The relative proportions of dense, lucent, and cargo-filled EVs differed significantly across classes (Figure 4C). Among large basiconic sensilla—ab1 (four ORNs), ab2 (two ORNs), and ab3 (two ORNs)—ab1 showed the greatest enrichment of cargo-filled EVs. Consistent with the larger size of cargo-filled EVs, ab1 sensilla also contained the highest proportion of EVs exceeding 200 nm (Figure 4D), consistent with a microvesicle-like size range (Malkin & Bratman, 2020; Shahi et al., 2024). In contrast, coeloconic and trichoid sensilla exhibited the highest proportions of dense EVs (Figure 4C).

Beyond density differences, we identified structural variants of EVs, including double EVs and ball-and-socket EVs. Double EVs, defined as vesicles that enclose one or more smaller vesicles (Figure 4E, left), have been described in human secretions and may permit protected cargo delivery by shielding their contents from lysosomal degradation in the recipient cytoplasm (Neyroud et al., 2022; Petersen et al., 2023; Saadeldin et al., 2023; Zabeo et al., 2017). The luminal densities of the inner and outer compartments were typically similar, though occasional variations occurred.

In contrast, ball-and-socket EVs consist of two vesicles physically coupled in a joint-like configuration, with one vesicle partially invaginated into the other (Figure 4E, right). These structures are predominantly >200 nm and typically located distally in the sensillum lumen, distant from the plasma membranes of ORNs or auxiliary cells where EV budding would occur. The “ball” and “socket” typically share similar densities (e.g., both lucent or both dense), though density variation is not a defining feature of this class. Unlike double EVs, which comprise a small intraluminal vesicle enclosed within a substantially larger outer vesicle, ball-and-socket configurations involve two similarly sized vesicles. Their size symmetry and distance from cell membranes suggest they represent physical interactions between mature extracellular vesicles rather than budding intermediates. To our knowledge, this morphology has not been previously described and may represent a distinct class of EVs or a previously unrecognized form of inter-vesicular interaction.

To determine how these structural variants relate to density categories, we quantified their composition and found that both were highly enriched in the cargo-filled category (Figure 4F), indicating a close association between complex EV morphology and cargo-rich internal structure. Consistent with this relationship, a large fraction of double and ball-and-socket EVs fall within the cargo-filled population, although each class also contained unique subsets (Figure 4G). Together, these findings demonstrate that EVs in the sensillum lymph are heterogeneous along multiple parameters—including size, intraluminal density, and structure—and that these properties are differentially distributed across sensillum classes.

### Multivesicular bodies within auxiliary cells

Because multivesicular bodies (MVBs)—endosomal compartments containing intraluminal vesicles—are key intracellular sources of exosomes (Doyle & Wang, 2019; Malkin & Bratman, 2020; Neyroud et al., 2022; Remans et al., 2025; Welsh et al., 2024), we examined their presence and distribution. Among sensilla where ORNs and all three auxiliary cell types were sampled (*n* = 304), 53 sensilla (∼17%) contained at least one MVB-positive auxiliary cell, whereas none were detected in ORNs, indicating that MVBs are restricted to auxiliary cells. Of note, MVBs were observed in all three auxiliary cell types: thecogen, trichogen, and tormogen cells (Figure 5A–C).

**Figure 5.**
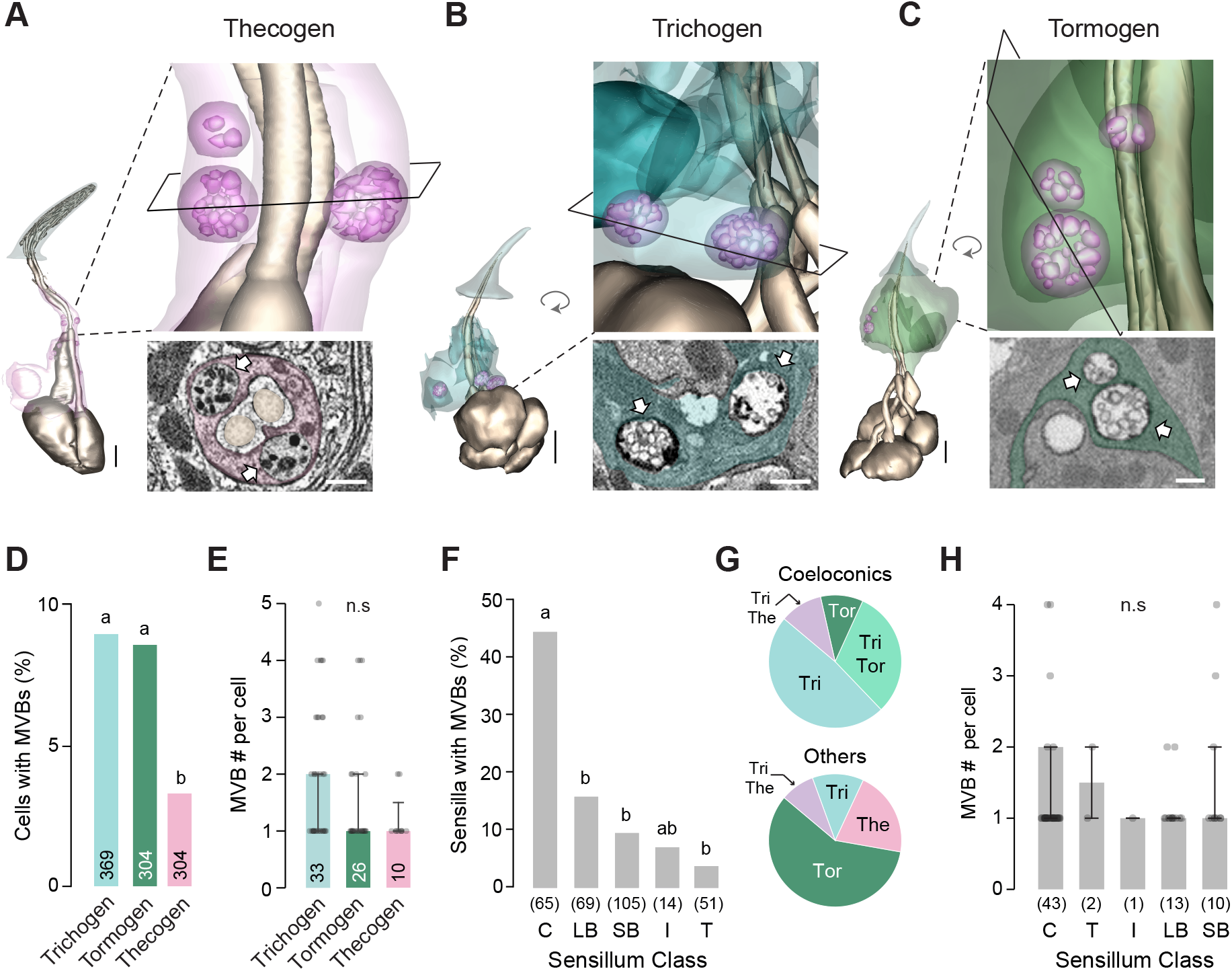
Multivesicular bodies within auxiliary cells. **(A)** 3D reconstructions of the sensillum cuticle (gray), ORNs (bronze), thecogen cell (light pink), and MVBs (dark pink). Inset: enlarged view showing the cutting plane for the representative SBEM image. MVBs are marked by thick white arrows. Scale bars: 2 µm (3D models) and 1 µm (SBEM image). (**B□C**) Same as (**A**), except different auxiliary cells are shown: (**B**) trichogen cell (turquoise) and (**C**) tormogen cell (green). **(D)** Percentage of auxiliary cells containing MVBs, observed more frequently in trichogen and tormogen cells. Sample sizes for each auxiliary cell type are indicated (from 304 sensilla). The higher number of trichogen cells reflects the presence of two trichogen cells per coeloconic sensillum, compared to one per sensillum in other sensillum types. Different letters denote statistically significant differences between groups (Chi-square test for independence with FDR-corrected Chi-square *post hoc* tests, *p* < 0.05). **(E)** Number of MVBs per auxiliary cell among cells containing at least one MVB. Sample sizes indicate the number of auxiliary cells with at least one MVB. Data are shown as median ± 95% CI. Kruskal–Wallis test, *p* > 0.05. **(F)** Percentage of sensilla containing MVBs (*n* = 304). MVBs were more frequently observed in coeloconic sensilla. Sample sizes indicate the number of surveyed sensilla. Statistical analysis as in (**D**). **(G)** Pie charts showing the distribution of auxiliary cell types containing MVBs in coeloconic snsilla (top, *n* = 29) and in all other sensillum types combined (bottom, *n* = 24). **(H)** Number of MVBs per auxiliary cell (cells with ≥1 MVB) across sensillum types. Median ± 95% CI. Sample sizes indicate the number of auxiliary cell with at least one MVB. Statistical analysis as in (**E**).

We first compared the prevalence of MVBs across auxiliary cell types. The percentage of cells containing MVBs differed significantly, with trichogen and tormogen cells showing higher prevalence than thecogen cells (Figure 5D). In contrast, among cells that contained at least one MVB, the number of MVBs per cell did not differ significantly across auxiliary cell types, with an overall median of 1 (Figure 5E).

We next examined MVB distribution across sensillum classes. Coeloconic sensilla exhibited the highest prevalence of MVB-containing auxiliary cells, accounting for 29 of the 53 MVB-positive sensilla (∼55%) from a total of 65 surveyed coeloconic sensilla (∼45%; Figure 5F). Within coeloconic sensilla, trichogen cells were the most common MVB-containing cell type, and in some cases, multiple auxiliary cell types within the same sensillum contained MVBs (Figure 5G, top). In contrast, among all other sensillum types combined, tormogen cells were the predominant MVB-containing cell type (Figure 5G, bottom).

The enrichment of MVBs in coeloconic sensilla was unexpected given our earlier observation that coeloconic sensilla exhibited the lowest proportion of EVs within the sensillum lumen (Figure 3C). One possibility is that coeloconic MVBs release exosomes into extracellular compartments outside the sensillum lumen. Alternatively, exosomes released into the coeloconic lumen may persist only briefly and therefore fail to accumulate to detectable levels.

Despite these differences in distribution, the number of MVBs per auxiliary cell was similar across sensillum classes, with medians ranging from 1 to 2 (Figure 5H). Together, these results indicate that while MVB occurrence varies across both auxiliary cell types and sensillum classes, their abundance within individual cells is relatively low, suggesting cell type- and sensillum-specific regulation of MVB presence.

### Putative EV budding from auxiliary cells

In addition to endosomal pathways mediated by MVBs, EVs—particularly microvesicles—can arise directly from the plasma membrane through outward budding (Remans et al., 2025; Singh et al., 2024). Across the same dataset of 304 sensilla, we identified 28 sensilla (∼9%) containing putative budding EVs, defined as vesicle-like structures whose membranes remained physically continuous with the plasma membrane of a parent cell (Figure 6). Because these observations derive from static ultrastructural images, we cannot distinguish vesicle budding from fusion events, and thus refer to them as putative budding EVs. All such structures were associated with auxiliary cells, with no instances detected in ORNs. Their occurrence was lower than that of MVB-containing sensilla (∼17%; Figure 5).

**Figure 6.**
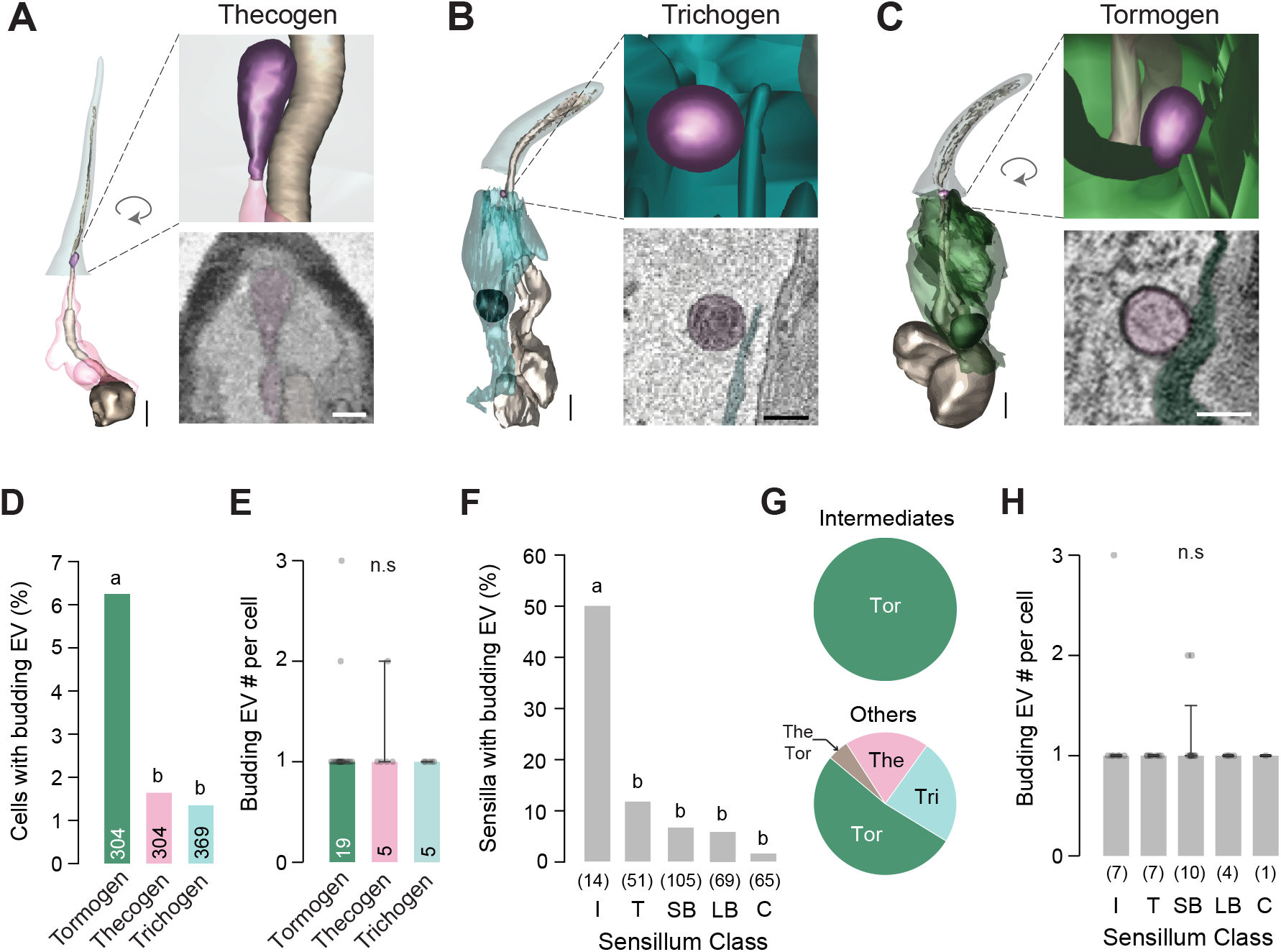
Putative EV budding from auxiliary cells. **(A)** 3D reconstructions of the sensillum cuticle (gray), ORNs (bronze), thecogen cell (light pink), and a putative budding EV (purple) that remains physically connected to the auxiliary cell membrane. Inset: enlarged view. Representative SBEM image is shown below. Scale bars: 2 µm (3D models) and 400 nm (SBEM image). (**B□C**) Same as (**A**), except different auxiliary cells are shown: (**B**) trichogen cell (turquoise) and (**C**) tormogen cell (green). In (**B**), the section shown captures a plane in which the EV membrane is no longer continuous with the trichogen cell membrane; membrane continuity is visible in adjacent serial sections (Figure 6—figure supplement 1). **(D)** Percentage of auxiliary cells with putative budding EVs, observed more frequently in tormogen cells. Sample sizes for each auxiliary cell type are indicated (from 304 sensilla). Different letters denote statistically significant differences between groups (Chi-square test for independence with FDR-corrected Chi-square *post hoc* tests, *p* < 0.05). **(E)** Number of budding EVs per auxiliary cell among cells with at least one putative budding EV. Sample sizes indicate the number of auxiliary cells with at least one putative budding EV. Data are shown as median ± 95% CI. Kruskal–Wallis test, *p* > 0.05. **(F)** Percentage of sensilla containing putative budding EVs (*n* = 304), which were more frequently observed in intermediate sensilla. Sample sizes indicate the number of surveyed sensilla. Statistical analysis as in (**D**). **(G)** Distribution of auxiliary cell types associated with putative budding EVs in intermediate sensilla (top, *n* = 7) and all other sensillum types combined (bottom, *n* = 21). **(H)** Number of putative budding EVs per auxiliary cell (cells with ≥1 budding EV) across sensillum types. Median ± 95% CI. Sample sizes indicate the number of auxiliary cell with at least one putative budding EV. Statistical analysis as in (**E**).

Putative budding EVs were observed in all three auxiliary cell types (Figure 6A–C; Figure 6—figure supplement 1). These structures appeared as vesicle-like profiles that remained physically continuous with the plasma membrane of a parent cell. Their distribution was not uniform, with tormogen cells exhibiting the highest frequency of such events (Figure 6D). Despite differences in occurrence, the number of putative budding EVs per cell remained low and did not differ significantly across auxiliary cell types, with a median of one event per cell among cells exhibiting at least one membrane-associated vesicle (Figure 6E). Notably, these vesicles were relatively large, with a median diameter of 521 nm (*n* = 33), substantially exceeding that of the overall EV population (177 nm; Figure 2C) and falling within the size range commonly associated with microvesicles (Shahi et al., 2024).

Across sensillum classes, putative budding EVs were most frequently observed in intermediate sensilla, where 7 of 14 sensilla (∼50%) contained at least one such event (Figure 6F). Notably, in intermediate sensilla, all putative budding EVs were associated with tormogen cells (Figure 6G, top). In other sensillum types, tormogen cells remained the predominant source, followed by trichogen and thecogen cells (Figure 6G, bottom). However, the number of putative budding EVs per auxiliary cell remained consistently low across sensillum classes, with an overall median of one and no significant differences across groups (Figure 6H).

Overall, both MVBs and putative budding events were infrequent, with budding observed less often than MVBs, likely reflecting the highly transient nature of membrane budding compared to the longer intracellular persistence of MVBs (Corrigan et al., 2014; M. Sun et al., 2021). These findings suggest that EVs in olfactory sensilla could arise from both endosomal (MVB-mediated) and plasma membrane budding pathways in all three auxiliary cell types. However, the observed frequencies likely underestimate their true occurrence, as these are dynamic cellular processes and our SBEM analysis provides only a static snapshot of adult tissues.

### Degenerating ORNs across sensillum classes

In addition to sensilla with intact ORNs, we identified 32 sensilla containing ORNs with morphological features of apoptosis (Figure 7). Consistent with previous observations of ORN apoptosis (Chihara et al., 2014; Fernández-Hernández et al., 2020; Takeuchi et al., 2022), affected dendrites exhibited hallmark features of apoptotic degeneration, including truncation, fragmentation, and membrane blebbing (Nagata, 2018; Xu et al., 2019). These abnormalities were observed across sensillum classes, including 6 coeloconic, 10 small basiconic, 7 large basiconic, 1 intermediate, and 8 trichoid sensilla. In some sensilla, both ORNs appeared affected, whereas in others only a single ORN exhibited signs of degeneration. Together, these sensilla accounted for 47 affected ORNs out of 802 surveyed (∼6%). The observation of degenerating ORNs in these SBEM volumes provided a unique opportunity to examine the ultrastructure of apoptotic bodies and degeneration-associated extracellular particles *in situ* within native olfactory sensilla.

**Figure 7.**
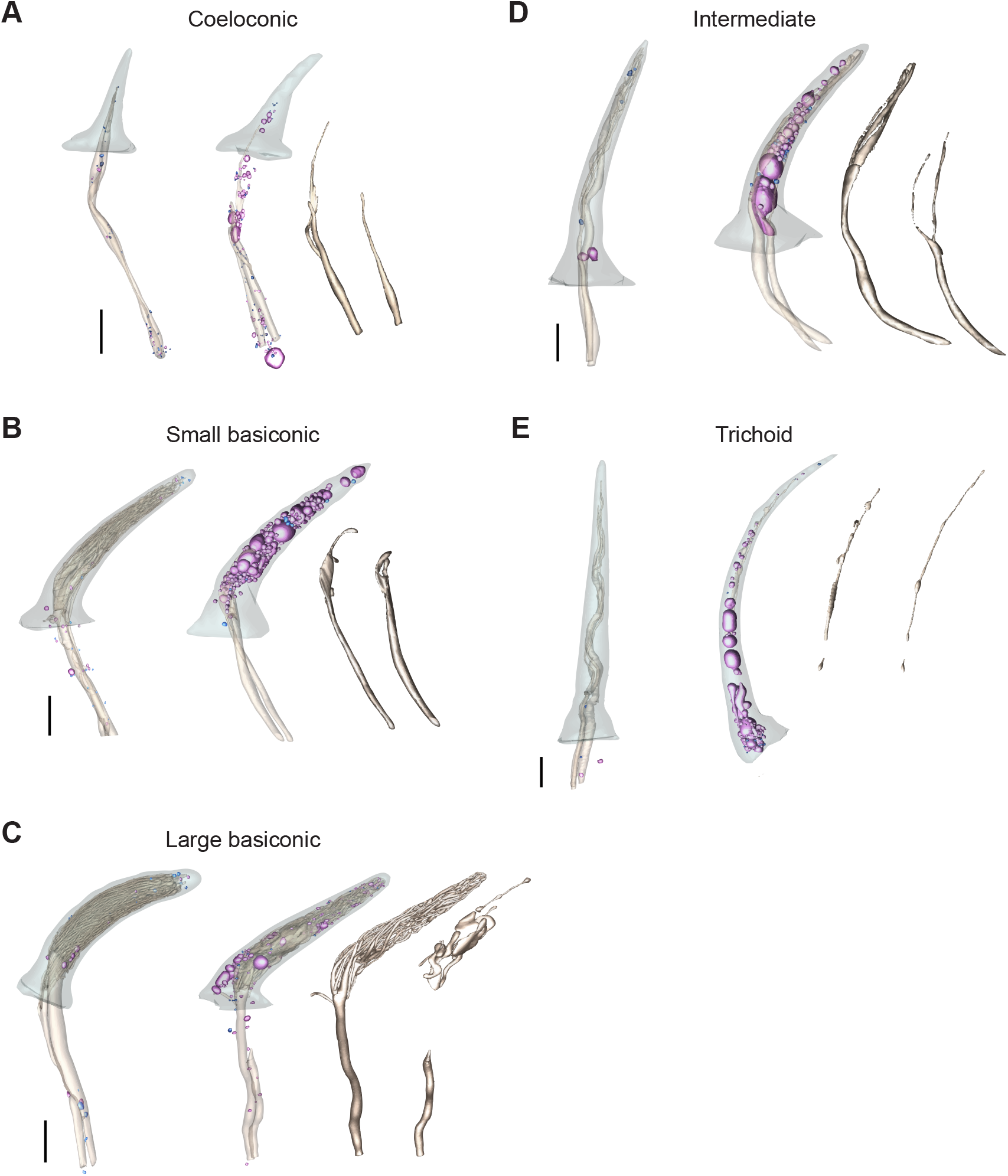
Degenerating ORNs across sensillum classes. **(A)** Representative 3D models of coeloconic sensillum showing the sensillum cuticle, ORN outer dendrites, EVs (purple), and NVEPs (blue; generally much smaller than EVs). Left: control sensillum with intact ORNs. Right: sensillum containing degenerating ORNs. The outer dendrites of both ORNs are truncated distally, and EVs and NVEPs are both more abundant relative to the control sensillum. Scale bar, 2 µm. **(B□E)** Same as (**A**), showing additional sensillum classes. (**B**) Small basiconic sensillum, with both ORN outer dendrites truncated distally. (**C**) Large basiconic sensillum, with one ORN outer dendrite fragmented in the middle region and exhibiting distal blebbing, defined as the presence of rounded membrane protrusions along the dendritic shaft. (**D**) Intermediate sensillum, with one ORN outer dendrite truncated distally. (**E**) Trichoid sensillum, with both ORN outer dendrites truncated proximally.

We note that this survey was not intended to provide an exhaustive quantification of ORN degeneration or to estimate the frequency of degenerating ORNs. Our flies carry the *UAS-APEX2* transgene, and even low-level leakage expression of this peroxidase might elevate reactive oxygen species and thereby increase susceptibility to apoptosis; therefore, these frequencies should be interpreted with caution. Rather, these observations demonstrate that ORN degeneration can occur across multiple sensillum classes and provided a unique opportunity to compare the morphological properties of EVs and NVEPs in sensilla with either degenerating or intact ORNs. Hereafter, sensilla containing intact ORNs are referred to as “intact sensilla”, whereas sensilla containing degenerating ORNs are referred to as “degenerating sensilla”.

### Comparative analysis of intact and degenerating sensilla

#### EV and NVEP number

Degenerating sensilla exhibited a marked increase in extracellular particles within the sensillum lumen (Figure 7). Across the 32 sensilla containing degenerating ORNs, we identified 3,396 EVs and 1,116 NVEPs, compared to 1,397 EVs and 1,914 NVEPs across 320 intact sensilla. The median number of NVEPs per sensillum was approximately fourfold higher in degenerating sensilla than intact sensilla (23.5 vs. 5), whereas the median number of EVs per sensillum was over 32-fold higher (100.5 vs. 3; Figure 8A). Thus, both EVs and NVEPs were enriched in degenerating sensilla, with EVs exhibiting particularly pronounced accumulation.

**Figure 8.**
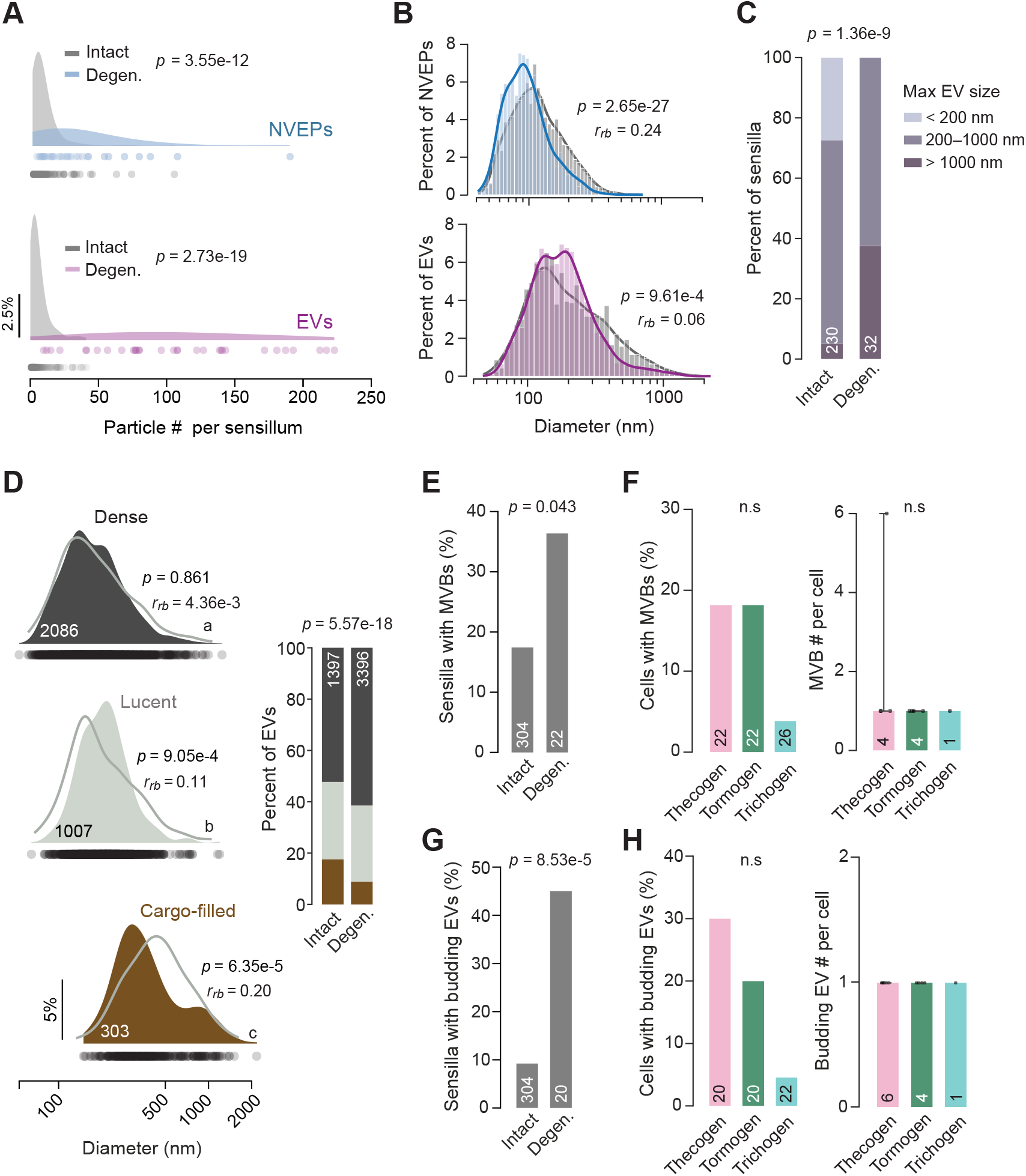
Comparison of extracellular particles between intact and degenerating sensilla. **(A)** EV and NVEP abundance per sensillum. Distributions are shown as kernel density estimates with individual data points plotted along the x-axis. Comparisons were made between intact sensilla (gray; *n* = 257 for NVEPs, *n* = 230 for EVs) and degenerating sensilla (colored; *n* = 32 for both) using Mann–Whitney U tests. Scale bar, 2.5% density. **(B)** Distributions of EV and NVEP diameters. Curves represent kernel density estimates. Differences between intact (gray) and degenerating sensilla (colored) were assessed using Mann–Whitney U tests, with effect sizes quantified by rank-biserial correlation (*r_rb_*) (*n* = 3,396 EVs; *n* = 1,116 NVEPs). Intact sensilla data are the same as in Figure 2C. **(C)** EV diameter categories. Stacked bar plots show proportions of maximum EV size categories in intact and degenerating sensilla. EVs >1000 nm fall within an apoptotic body-like size range. Statistical comparison was performed using Chi-square tests for independence. **(D)** Size distributions of EVs grouped by intraluminal density (degenerating sensilla). Gray line: distribution for intact sensilla (as in Figure 4B). When comparing intact vs degenerating sensilla within each intraluminal density, statistical analysis is as in (**B**). When comparing between intraluminal density categories within degenerating sensilla, different letters denote statistically significant differences between groups (Kruskal–Wallis test with FDR-corrected Dunn’s *post hoc* comparisons, *p* < 0.05). Hartigan’s dip test supported unimodality for dense (*p* = 0.994), lucent (*p* = 0.964), and cargo-filled EVs (*p* = 0.995). Inset: Stacked bar plots showing the distribution of intraluminal density categories between intact and degenerating sensilla, highlighting enrichment of the dense EV class in degenerating sensilla. Statistical analysis as in (**C**). **(E)** Percentage of sensilla containing auxiliary cells with MVBs. Only sensilla with intact auxiliary cells were included. Statistical comparison was performed using Fischer exact test. **(F)** Left: Percentage of auxiliary cells containing MVBs in degenerating sensilla. Chi-square test for independence, *p* > 0.05. Right: Number of MVBs per auxiliary cell, shown as median ± 95% CI. Kruskal–Wallis test, *p* > 0.05. (**G□H**) Same as (**E□F**), except analyses were performed for putative budding EVs. Statistical comparisons could not be performed for the right panel of (**H**) because all auxiliary cells contained a single budding EV.

#### EV and NVEP size and position

Similar to our observations in intact sensilla (Figure 2C), NVEPs were generally smaller than EVs in degenerating sensilla. Analysis of continuous size distribution showed that the median diameters of NVEPs and EVs were 92 nm and 173 nm, respectively (Figure 8B). When comparing intact and degenerating sensilla, both NVEPs and EVs were modestly but significantly smaller in degenerating sensilla at the population level.

However, a striking pattern emerged when sensilla were categorized by the size of their largest EVs. Approximately 27% of intact sensilla only contained small EVs (<200 nm, the likely size range of exosomes), whereas this category was absent in degenerating sensilla (Figure 8C). Conversely, only ∼5% of intact sensilla contained very large EVs (>1000 nm, large EVs and apoptotic bodies), compared to ∼38% of degenerating sensilla.

The categorical analysis reveals an additional, biologically relevant pattern that the continuous distribution alone can obscure: degeneration-associated extracellular particle accumulation is primarily driven by increased particle abundance across all size ranges. Notably, all degenerating sensilla contained EVs >200 nm, and a substantially larger fraction contained very large EVs (>1000 nm). Although apoptotic bodies are highly heterogeneous in size and can range broadly from ∼50 nm to several microns (Elsner et al., 2023), these large EVs fall within the upper size range commonly associated with apoptotic bodies (Battistelli & Falcieri, 2021; Shahi et al., 2024).

Furthermore, the spatial distributions of both particle types were shifted modestly toward the distal end of the sensillum lumen, resulting in median positions that were slightly farther above the cuticle base than those observed in intact sensilla (Figure 8—figure supplement 1). Because the lumen above the cuticle base contains ORN outer dendrites, this subtle distal shift suggests that degeneration-associated particle accumulation may occur preferentially within the dendrite-containing region of the sensillum.

#### Intraluminal electron density

We next examined whether EV intraluminal electron density profiles are altered in degenerating sensilla. Similar to intact sensilla, EVs in degenerating sensilla could also be classified into three categories by their intraluminal density: dense, lucent, and cargo-filled (Figure 8D). Cargo-filled EVs remained the category with the largest diameters and exhibited a broad right-skewed size distribution (median diameter = 339 nm), followed by lucent and dense EVs (185 nm and 149 nm, respectively). Thus, the overall relationship between EV intraluminal density and vesicle size was preserved in degenerating sensilla.

Notable differences emerged when intact and degenerating sensilla were compared across vesicle density categories (Figure 8D). Lucent EVs were larger in degenerating sensilla than in intact sensilla (185 vs. 161 nm), whereas cargo-filled EVs were markedly smaller (339 vs. 440 nm). In addition, degenerating sensilla contained a higher proportion of dense EVs (∼61%) relative to intact sensilla (∼52%), accompanied by a reduction in cargo-filled EVs (∼9% vs. ∼18% in intact sensilla). These findings suggest that ORN degeneration is associated not only with increased EV accumulation, but also with shifts in EV composition and size distribution. The increased abundance of dense EVs, which contain high concentrations of biomolecular material (Shahi et al., 2024), may partially reflect the presence of apoptotic bodies.

#### Multivesicular bodies and putative budding EVs

To evaluate EV biogenesis pathways in degenerating sensilla, we quantified the occurrence and cellular distribution of MVBs and putative budding EVs. Analyses were restricted to sensilla with intact auxiliary cells for MVB quantification (*n* = 22), and to sensilla with intact auxiliary cells exhibiting well-preserved microlamellar morphology for budding EV analysis (*n* = 20).

MVB-containing auxiliary cells were identified in ∼36% of degenerating sensilla, representing a significant increase compared to ∼17% of intact sensilla (Figure 8E). The distribution of MVBs across auxiliary cell types also differed between the two groups. In degenerating sensilla, MVBs were more frequently observed in thecogen and tormogen cells, although these differences were not significant (Figure 8F, left). In contrast, intact sensilla showed higher MVB occurrence in trichogen and tormogen cells than in thecogen cells (Figure 5D). Despite these differences, the median number of MVBs per auxiliary cell remained at one across all cell types, similar to intact sensilla (Figure 8F, right).

We next analyzed budding events associated with EV release. Auxiliary cells containing putative budding EVs were identified in ∼9% of intact sensilla and ∼45% of degenerating sensilla, indicating a marked increase in degenerating sensilla (Figure 8G). In degenerating sensilla, putative budding EVs were most frequently observed in thecogen cells, followed by tormogen and trichogen cells, although these differences were not significant (Figure 8H, left). This distribution differed from intact sensilla in which putative budding events were most common in tormogen cells (Figure 6D). Similar to MVBs, the median number of putative budding EVs per auxiliary cell remained one regardless of cell type (Figure 8H, right).

Together, these findings suggest that ORN degeneration is associated with shifted auxiliary cell involvement in EV biogenesis pathways, specifically via greater participation of thecogen cells. However, because degenerating ORNs themselves exhibited extensive dendritic fragmentation and blebbing, a substantial fraction of extracellular particles is expected to originate directly from degenerating ORNs as apoptotic bodies. Thus, the increased EV abundance in degenerating sensilla likely reflects contributions from both enhanced auxiliary cell EV biogenesis and degenerating ORNs themselves.

### Degenerating auxiliary cells

In addition to sensilla containing degenerating ORNs, we identified a second class of degenerating sensilla in which the ORNs remained morphologically intact, while the associated auxiliary cells exhibited disintegrated plasma membranes (Figure 9A and B). These auxiliary cells displayed morphological features more consistent with necrotic/necroptotic degeneration, unlike apoptotic ORNs, which showed dendritic blebbing while maintaining intact plasma membranes (Dhuriya & Sharma, 2018; Smith et al., 2024; Ye et al., 2023).

**Figure 9.**
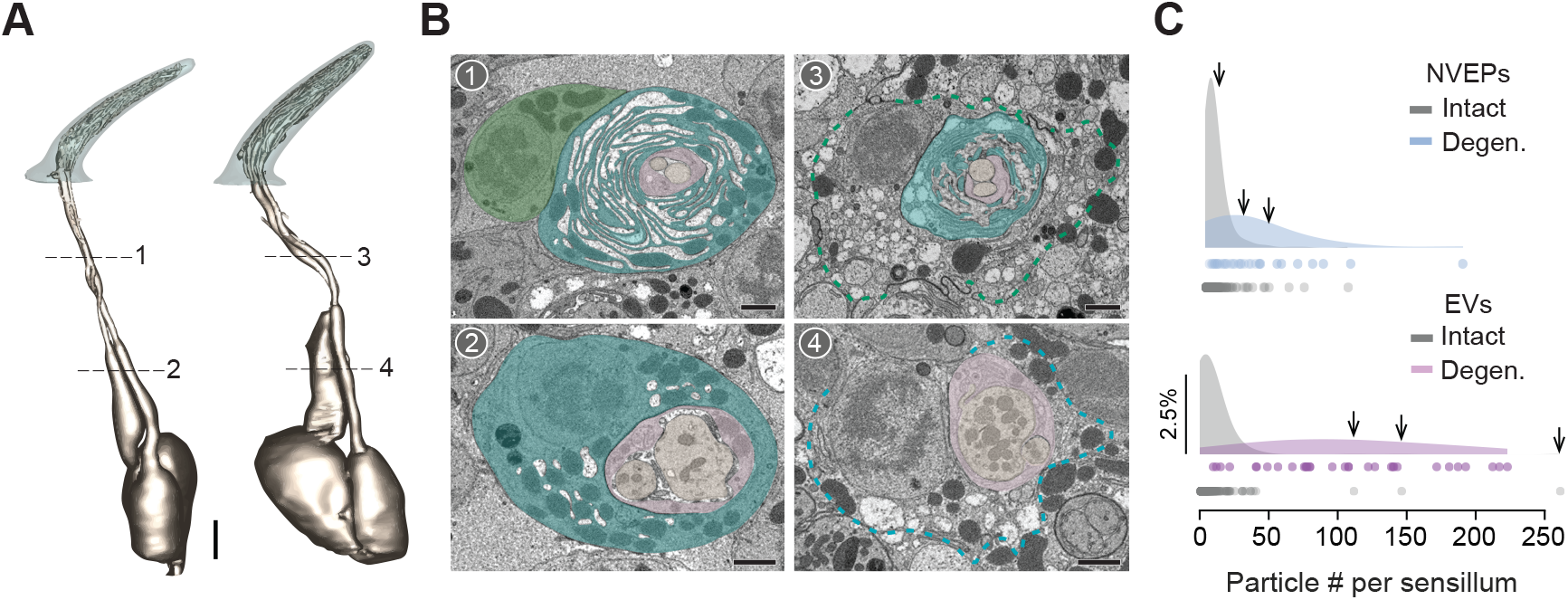
Sensilla containing degenerating auxiliary cells but intact ORNs. (**A□B**) Representative 3D models (**A**) and corresponding SBEM images (**B**) of two basiconic sensilla from the same SBEM volume, showing the sensillum cuticle and intact ORNs. Left: control sensillum with intact auxiliary cells. Right: sensillum containing degenerating auxiliary cells. Cells are pseudocolored by identity: tormogen cell (green), trichogen cell (turquoise), thecogen cell (pink), and ORNs (bronze). Dashed outlines in images 3 and 4 indicate disintegrated membranes of the tormogen and trichogen cells, respectively. Scale bars, 2 µm for 3D models and 1 µm for SBEM images. (**C**) EV and NVEP abundance per sensillum, as in Figure 8A. Arrows indicate datapoints from the three sensilla containing degenerating auxiliary cells but intact ORNs.

We analyzed three sensilla of this type and found that they also contained elevated numbers of extracellular particles within the sensillum lumen—particularly EVs—similar to sensilla with degenerating ORNs (Figure 9C). These observations suggest that degeneration-associated extracellular particles are not exclusively produced by apoptotic ORNs, but also arise in response to auxiliary cell degeneration. The increased abundance of extracellular particles in sensilla with degenerating auxiliary cells further raises the possibility that degeneration-associated signals released from auxiliary cells may be communicated to ORNs through NVEPs and EVs, to thereby promote ORN degeneration.

Indeed, 5 of the 32 sensilla containing degenerating ORNs also exhibited degenerating auxiliary cells (Figure 9—figure supplement 1), suggesting that degeneration can initiate from either ORNs or auxiliary cells. However, degeneration appears to ultimately spread to all cellular components within the sensillum, consistent with our published observation of “empty sensilla” in which both ORNs and auxiliary cells had degenerated within the sensillum cuticle (Nava Gonzales et al., 2021).

## DISCUSSION

Here, we provided a systematic ultrastructural survey of extracellular particles within olfactory sensilla and showed that both EVs and NVEPs are widespread but heterogeneously distributed across sensillum classes. By preserving the native cellular context through cryofixation-based volume EM, our analyses uncovered distinct particle populations, sensillum class-specific distributions, and ultrastructural signatures of EV biogenesis that would be difficult to resolve with conventional light microscopy or dissociated preparations.

Our findings also highlight the substantial heterogeneity across olfactory sensilla. Individual sensillum classes house distinct ORNs tuned to specific odors that regulate diverse behaviors, including foraging, egg laying, avoidance, and courtship (Wu et al., 2022). Accordingly, the extracellular environments surrounding these neurons are also likely to be functionally specialized. For example, ab1—a large basiconic sensillum uniquely containing four ORNs, including the CO_2_-sensing neuron (Choy et al., 2025; de Bruyne et al., 2001)—selectively enriches large, cargo-filled vesicles (Figure 4C), suggesting EV-mediated communication tailored to the demands of this sensillum population. This heterogeneity should therefore be carefully considered in future studies of EVs and NVEPs in sensory signaling, as mechanisms identified in one sensillum class may not generalize across the olfactory system. Rather than a one-size-fits-all model, extracellular particle signaling in sensory tissues is likely shaped by the unique physiological and behavioral roles of individual olfactory processing units.

The anatomical organization of olfactory sensilla further suggests that these particles primarily mediate local intercellular communication within individual sensilla. The vast majority of EVs and NVEPs observed in this study greatly exceeded the diameter of cuticular pores, which are typically ≤10 nm in diameter (Shanbhag et al., 1999). Consequently, unlike extracellular vesicles involved in animal-to-animal communication in *C. elegans* (Wang et al., 2014), extracellular particles within *Drosophila* olfactory sensilla are unlikely to escape into the external environment. Instead, they are positioned to influence interactions among ORNs and auxiliary cells within the enclosed sensillum lumen, where they may contribute to the local regulation of sensory physiology and signaling.

EVs are thought to arise through endosomal release via MVBs or direct plasma membrane budding (Doyle & Wang, 2019; Malkin & Bratman, 2020; Neyroud et al., 2022; Remans et al., 2025; Singh et al., 2024; Welsh et al., 2024). Our data suggest both pathways are active but differentially favored across sensillum classes. Of note, coeloconic sensilla contain abundant small, dense EVs alongside frequent intracellular MVBs (Figures 4C and 5F), consistent with endosomal release. Conversely, ab1 sensilla are enriched in large, cargo-filled vesicles with fewer MVBs, suggesting a preference for direct plasma membrane budding. The scarcity of both MVBs and budding figures in intact sensilla likely reflects the transient nature of EV biogenesis.

Ciliary extracellular vesicle shedding is well documented across evolutionarily distant systems, from the flagella of *Chlamydomonas* to the sensory cilia of *C. elegans* and mammalian primary cilia (Wang & Barr, 2016, 2018). It is therefore notable that we did not capture active EV budding from ORN dendritic cilia in our adult *Drosophila* preparations. This absence likely reflects both biological and technical constraints: ciliary EV production may be developmentally regulated and reduced in 1-week-old adults relative to earlier stages, and static volume EM may miss rare, transient release events. Instead, our data suggest that auxiliary cells are active contributors to the luminal EV pool in adults, a view independently supported by the recent identification of support-cell-specific *GAL4* drivers that label EV-like structures originating from tormogen and trichogen membranes (Ballou et al., 2026). EV-mediated exchange between sensory neurons and supporting cells appears to be evolutionarily conserved. In *Drosophila* olfactory sensilla, where ORNs are intimately ensheathed by auxiliary cells, EV-mediated crosstalk may similarly coordinate sensillar physiology, though future functional studies with cell-specific genetic manipulation will be needed to test this directly.

The marked increase in extracellular particles within degenerating sensilla initially suggested that these structures arise directly from apoptotic ORNs. Indeed, degenerating ORNs exhibited classic hallmarks of apoptotic degeneration, including dendritic truncation, fragmentation, and blebbing (Nagata, 2018; Xu et al., 2019), and many of the large dense EVs observed in degenerating sensilla fell within the size range associated with apoptotic bodies (Figure 8C). However, our analyses also revealed elevated EV biogenesis in auxiliary cells, as well as shifts in the auxiliary cell types participating in MVB- or budding-associated EV production (Figure 8E H). That is, the elevated EV abundance in degenerating sensilla originates from multiple cellular sources, rather than exclusively from degenerating ORNs themselves. The increased involvement of thecogen cells is particularly intriguing given that they closely envelop ORNs, suggesting that they may respond directly to signals of neuronal distress. Furthermore, the shift toward dense EVs in degenerating sensilla indicates altered extracellular environment (Figure 8D). These particles may represent increased apoptotic bodies containing condensed cellular material, as well as stress-induced vesicles released by auxiliary cells.

Our findings raise the intriguing possibility that auxiliary cells actively participate in degeneration within olfactory sensilla. In previous studies, we identified rare “empty sensilla” lacking all cellular components (Nava Gonzales et al., 2021), suggesting that degeneration can ultimately lead to coordinated loss of all cells within an individual sensillum. The sparse and isolated distribution of empty sensilla further suggests that degeneration remains spatially restricted. Importantly, our observation of auxiliary cell degeneration with intact ORNs—and vice versa—indicates that degeneration can be initiated in either cellular component and subsequently spread to other cells within the same sensillum over time, revealing coordinated multicellular remodeling during sensory degeneration. Further molecular studies will be required to determine whether extracellular vesicles directly contribute to degeneration-associated intercellular signaling within olfactory sensilla.

An important unanswered question concerns the lifetime and turnover of extracellular particles within olfactory sensilla. Although our analyses identify their abundance, distribution, and potential cellular origins, little is known about how long these particles persist or how they are ultimately cleared from the sensillum lumen. Notably, despite the frequent presence of EVs and NVEPs in the dendrite-containing lumen above the cuticle base, we did not observe convincing ultrastructural evidence of particle fusion with or endocytic uptake by ORN dendrites. Future studies examining extracellular particle turnover and degradation will be important for understanding how sensory tissues regulate intercellular signaling.

## Materials and Methods

### SBEM volumes

The SBEM volumes analyzed in this study were generated previously (Choy et al., 2025; Nava Gonzales et al., 2021). Briefly, 6- to 8-day-old female flies were cold-anesthetized, and their antennae were dissected. After isolation of the third antennal segment, samples were processed using high-pressure freezing, freeze substitution, rehydration, DAB labeling, *en bloc* heavy metal staining, dehydration, and resin infiltration. Flies expressed membrane-tethered APEX2 (*10xUAS-myc-APEX2-Orco* or *10xUAS-mCD8GFP-APEX2*) in select ORNs under the control of specific *OrX-GAL4* drivers, as described previously (Tsang et al., 2018; Zhang et al., 2019). Resin-embedded specimens were imaged by micro-computed X-ray tomography to determine sample position and orientation, then mounted on aluminum pins using conductive silver epoxy and sputter-coated with gold-palladium. SBEM imaging was performed using either a Gemini SEM (Zeiss) equipped with a 3View block-face unit or a Merlin SEM (Zeiss) equipped with a 3View2XP system and OnPoint backscatter detector. After data collection, the images were converted to MRC format, and rigid alignment of the image slices was performed using cross-correlation in the IMOD image processing package (https://bio3d.colorado.edu/imod/).

The SBEM image volumes are available in the Cell Image Library (http://www.cellimagelibrary.org/home) under the following accession numbers: CIL:57519 (Volume 1), CIL:54614 (Volume 2), CIL:54609 (Volume 3), CIL:57579 (Volume 4). The segmented 3D models (including EVs, NVEPs, ORNs, and sensilla) are available separately through the Dryad Digital Repository (DOI: 10.5061/dryad.3j9kd521p). Segmented structures were grouped according to their SBEM volume of origin (volumes 1–4) and organized into structural groups using the built-in object list dialog in IMOD (accessed via “Edit” and then “Object list” in the Model View window). This grouping allows users to extract morphometric information for selected structural groups. Detailed information about the object list function is provided in the IMOD user guide (https://bio3d.colorado.edu/imod/doc/3dmodHelp/objectList.html).

### Image segmentation

Manual segmentation was performed in IMOD using drawing tools to place closed contours around structures of interest across serial sections (Kremer et al., 1996). ORNs and sensillum cuticles were segmented as described (Nava Gonzales et al., 2021). The ciliary constriction was used to define the boundary between inner and outer dendritic segments of ORNs (Shanbhag et al., 1999).

Extracellular vesicles (EVs) were identified by the presence of a clearly discernible membrane, appearing as a dark boundary similar to membranes of ORNs and support cells (Nava Gonzales et al., 2021; Shanbhag et al., 1999; Van Niel et al., 2018). Non-vesicular extracellular particles (NVEPs) were defined as extracellular structures lacking a clearly discernible membrane. Only EVs and NVEPs located in the sensillum lumen above the ciliary constriction were included in the analysis.

### Morphometric analysis

EVs, NVEPs, sensillum cuticle, ORN soma, and inner and outer dendritic segments were saved as separate objects to facilitate morphometric analysis. Segmented objects were subsequently meshed to connect adjacent contours and generate continuous 3D reconstructions. Detailed information about “imodmesh” and IMOD’s drawing tools is available in the IMOD user guide (https://bio3d.colorado.edu/imod/doc/man/imodmesh.html; https://bio3d.colorado.edu/imod/doc/3dmodHelp/plughelp/drawingtools.html).

Morphometric measurements were extracted from individual objects using the “imodinfo” function in IMOD. The “Volume Inside Mesh” option was used to quantify object volumes. Effective diameter of EVs and NVEPs (D, μm) was estimated using the formula 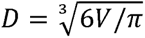 and sphericity Φ (unitless) was calculated as 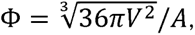 where *V* is volume (μm^3^) and *A* is surface area (μm^2^). Detailed documentation for “imodinfo” is available in the IMOD user guide (https://bio3d.colorado.edu/imod/doc/man/imodinfo.html).

To measure the absolute length of individual cuticle objects, each structure was first skeletonized in Amira (2020.2 version; ThermoFisher Scientific, USA) to generate a centerline represented as connected points. The resulting SWC files were imported into Python, where coordinates were converted from pixels to micrometers using scaling factors obtained from “imodinfo”. Segment lengths were calculated using the Pythagorean theorem, and the total cuticle length was obtained by summing all segments.

### Sensillum type identification

Sensilla were classified into major morphological types—large basiconic, small/thin basiconic, trichoid, intermediate, and coeloconic—based on cuticular morphology, including length, volume, and shape (Nava Gonzales et al., 2021). Large basiconic sensilla were further subdivided by neuronal content: ab1 sensilla contain four neurons, whereas ab2 and ab3 sensilla each contain two neurons. The thin (ab4) and small (ab5–ab11) basiconic sensilla could not be readily distinguished based on morphology and were therefore grouped together and referred to as small basiconic sensilla for simplicity. To distinguish intermediate sensilla from small/thin basiconic sensilla, isosurface models were generated and examined for diagnostic structural features. Although these sensillum types are similar in size, intermediate sensilla are defined by the presence of a basal drum structure, which is absent in basiconic sensilla (Nava Gonzales et al., 2021).

Isosurface models were generated using IMOD (Kremer et al., 1996) with the “Isosurface” function. Bounding box parameters were used to define regions of interest and extract subvolumes for rendering, and threshold values were manually adjusted based on image quality.

### Mapping of EV and NVEP positions along the sensillum cuticle

To determine the relative positions of EVs and NVEPs along the sensillum cuticle, cuticle skeletons in SWC format were imported into Python. Each EV or NVEP was projected onto the nearest point along the cuticle skeleton. Position was expressed as a normalized cuticle coordinate, scaled from 0 at the cuticle base to 1 at the cuticle tip. For particles located below the cuticle base, the terminal segment of the skeleton was extended, and positions were projected onto this extrapolated segment. In these cases, coordinates extended below 0, with −1 corresponding to one cuticle length below the base.

### Classification of EV luminal density and morphology

The inner luminal electron density of individual extracellular vesicles (EVs) was visually classified into three categories: dense, lucent, and cargo-filled. To account for variations in staining intensity and contrast across EM images, EV luminal density was evaluated relative to the surrounding lumen background of the same image. Dense EVs exhibited luminal density similar to or greater than that of the surrounding sensillum lumen, whereas lucent EVs displayed lower luminal density. Both dense and lucent EVs were defined by the uniformity of that density throughout the EV interior. On the other hand, cargo-filled EVs contained heterogeneous, electron-dense material within the lumen. In addition, distinct morphological configurations were annotated, including double EVs, defined as vesicles containing one or more internal vesicles, and ball-and-socket structures, in which a projection from one EV inserts into an invagination of another.

### Sensillum lumen volume estimate

Inner lumen volumes for the 352 sensilla analyzed in this study were estimated by multiplying each measured cuticle volume by a subtype-specific lumen-to-cuticle volume ratio (lumen volume / cuticle volume). These ratios were derived from a reference dataset of 150 genetically identified sensilla, in which the lumen and cuticle were segmented and their volumes were determined using the IMOD command *imodinfo* (Nava Gonzales et al., 2021). For coeloconic sensilla, which possess a double-walled cuticle (Shanbhag et al., 1999), the inner cuticular cylinder was used to define the lumen boundary. Lumen-to-cuticle volume ratios were calculated for each sensillum in the reference dataset, and subtype-specific mean ratios were subsequently used to estimate lumen volumes in the current study (see Source Data for Figure 3).

### SBEM image post-processing

For representative SBEM images of EVs and NVEPs, brightness and contrast were adjusted using the Contrast Limited Adaptive Histogram Equalization (CLAHE) function in FIJI/ImageJ. When necessary, images were additionally processed using the “Smooth” or “Sharpen” functions to enhance visual quality. These processing steps were applied for illustration purposes only and were not used for quantitative analyses.

### Outlier exclusion

For comparisons of EV and NVEP counts across sensillum classes (Figure 3D), outliers were identified within each sensillum class using Tukey’s extreme fences. Data points falling outside the range [*Q*1 - 3 × *IQR*, *Q*3 + *IQR*] were excluded. No outliers were removed in other analyses.

### Statistical analysis

Nonparametric statistical tests were used for the count, proportion, and morphological analyses. For comparisons of a continuous variable to a benchmark (e.g. comparing EV ratio to 0.5 for each sensillum class), a Wilcoxon signed-rank test was used. For comparisons of a continuous variable between two independent groups (e.g., EV vs NVEP sphericity), the Mann–Whitney U test was applied. If the Mann–Whitney U test yielded a significant result, the rank-biserial correlation (*r_rb_*) was calculated to quantify the effect size. For comparisons across three or more independent groups (e.g., EV number across sensillum morphological classes), the Kruskal–Wallis test was used. If the Kruskal–Wallis test yielded a significant result, Dunn’s post hoc pairwise comparisons were performed with Benjamini–Hochberg false discovery rate (FDR) correction. For categorical data (e.g., EV density distributions across sensillum classes), global differences were assessed using the Chi-square test for independence. If more than two groups were being compared and the global Chi-square test had a significant result, it was followed by pairwise Chi-square tests for independence with Benjamini–Hochberg FDR correction. If only two groups were being compared and there were only 2 categories (i.e. in Figure 8E and 8G), global differences were assessed using the Fischer exact test.

Statistical significance was defined as an adjusted *p*-value < 0.05. Differences among multiple groups were visualized using a compact letter display (CLD), in which groups sharing the same letter are not significantly different.

## Acknowledgments

We thank Joseph Luu and Melissa Zhao for assistance in EV segmentation, and Andreas Ernst, Andrew Chisholm, and Renny Ng for discussions and comments on the manuscript. This study was supported by NIH grants R01DC016466, R01DC021551 and R21DC020536 (C-Y.S.).

## Figure legends

**Figure 1—figure supplement 1. Schematic of a fly olfactory sensillum**

Two olfactory receptor neurons (ORNs) are housed within the sensillum; their dendrites extend into the sensillum lumen and are divided into inner and outer segments by the ciliary constriction. Three auxiliary cells—the thecogen (pink), trichogen (turquoise), and tormogen cells (green)—ensheathe the ORNs. The trichogen and tormogen cells extend microlamellae that line the bottom of the lumen, which is capped by the porous sensillum cuticle. Surrounding epithelial cells are shown in gray, and tight junctions between cells are represented by think horizontal lines.

**Figure 3—figure supplement 1. Relative positions of NVEPs and EVs along the sensillum axis.**

**(A)** Schematic illustrating particle positions relative to the cuticle base. The cuticle base was defined as 0, the sensillum tip as 1, and one sensillum cuticle length below the cuticle base as −1.

**(B–C)** Violin plots showing the relative positions of NVEPs (**B**) and EVs (**C**) along the sensillum axis. The numbers of particles and sensilla are listed in the Source Data for Figure 3S1. The sensillum classes are intermediate (I), small basiconic (SB), trichoid (T), large basiconic (LB), and coeloconic (C). Different letters indicate statistically significant differences among sensillum classes (Kruskal–Wallis test followed by FDR-corrected Dunn’s post hoc tests, *p* < 0.05).

**Figure 6—figure supplement 1. Serial SBEM images showing membrane continuity between putative budding EVs and auxiliary cells**

**(A)** Sample SBEM image of a putative EV (purple) budding from trichogen microlamellae (turquoise) in an ab1 sensillum. Magnified serial sections of the boxed region are shown on the right. Membrane continuity between the EV and trichogen cell membrane is visible in images 1 and 2, whereas images 3 and 4 show sections in which the EV membrane is separated from the trichogen cell membrane. Scale bars, 400 nm.

**(B)** Sample SBEM image of a putative budding EV (purple) from tormogen microlamellae (green) in an ai2 sensillum. Magnified serial sections of the boxed region are shown on the right. Membrane continuity between the EV and tormogen cell membrane is visible in image 3, whereas adjacent sections show the relative positions of the EV and tormogen cell in neighboring planes. Scale bars, 400 nm

**Figure 8—figure supplement 1. Comparison of NVEP and EV positions along the sensillum axis between intact and degenerating sensilla.**

**(A)** Schematic illustrating particle positions relative to the cuticle base. The cuticle base was defined as 0, the sensillum tip as 1, and one sensillum cuticle length below the cuticle base as −1.

(**B**–**C**) Violin plots showing the relative positions of NVEPs (**B**) and EVs (**C**) along the sensillum axis in intact and degenerating sensilla. The numbers of particles and sensilla are listed in the Source Data for Figure 8S1. Group differences were assessed using the Mann-Whitney U test, and effect sizes were quantified using the rank-biserial correlation (*r_rb_*).

**Figure 9—figure supplement 1. Sensilla containing degenerating auxiliary cells and ORNs**

(**A**) 3D models of ORNs housed in the same ab1 sensillum. The dendrite of “A” neuron shows blebbing (arrow heads) and fragmentation. Scale bar, 2 µm.

(**B**) Representative SBEM images of the sensillum. Cells are pseudocolored by identity: tormogen cell (green), trichogen cell (turquoise), thecogen cell (pink), and ORNs (bronze). Dashed outlines indicate disintegrated membranes of the trichogen cell (top) and tormogen cells (bottom). Scale bar, 1 µm.

